# A population genomics approach to unlock the genetic potential of lablab (*Lablab purpureus*), an underutilized tropical forage crop

**DOI:** 10.1101/2023.11.27.568768

**Authors:** A. Teshome, E. Habte, J. Cheema, A. Mekasha, H. Lire, M.S. Muktar, J. Quiroz-Chavez, C. Domoney, C.S. Jones

**Affiliations:** Feed and Forage Development, International Livestock Research Institute, Addis Ababa, Ethiopia; Ethiopian Institute of Agricultural Research Melkassa Agricultural Research Centre, Melkassa, Ethiopia; Ethiopian Institute of Agricultural Research Wondogenet Agricultural Research Centre, Wondogenet, Ethiopia; John Innes Centre, Norwich Research Park, Norwich NR4 7UZ, UK

**Keywords:** Dolichos, Genetic diversity, hyacinth bean, Genome-wide association study, Marker-assisted breeding

## Abstract

**Background:** In Sub-Saharan Africa (SSA) livestock production and productivity are severely restricted by the scarce supply of feedstuffs and forage crops, while those available are often of low nutritional quality resulting in poor animal productivity and leading to widespread malnutrition among the public, particularly women and children. Traditionally, several tropical forage crops have been used in the region both in the rangelands and as cut-and-carry cropping systems, but limited research attention has been paid to improve the quality of and access to these animal feeds. Lablab (*Lablab purpureus* (L.) is one of the conventionally grown multi-purpose underutilized crops that originated in Africa. It is an annual or short-lived perennial multi-purpose forage legume which has versatile uses (as a vegetable and dry seed), and as feed for animals, or as green manure. To develop new and highly productive lablab varieties, using genomics-assisted selection, the present study aimed to identify quantitative trait loci (QTL) associated with agronomically important traits in lablab and to assess the stability of these traits across two different agro-ecologies in Ethiopia.

**Results:** One hundred and forty-two cultivated and wild lablab accessions displayed significant agro-morphological variation in eight analysed traits, including plant height, total fresh weight, and total dry weight. Further, the agronomic performance of the accessions was significantly different across locations and years, highlighting the substantial genotype-by-environment interactions. The population genetic structure of the lablab accessions, based on half a million high quality single nucleotide polymorphisms (SNPs), revealed an independent domestication pattern for two-seeded and four-seeded lablab accessions. Furthermore, based on multi-environmental trial data, a genome-wide association study (GWAS) identified useful SNPs and *k*-mers for yield-related traits, such as plant height and total dry weight.

**Conclusions:** Genomic-assisted breeding is playing a key role in accelerating trait improvement in temperate forages, such as perennial ryegrass and alfalfa. Here we show that a similar approach could benefit underutilized crops such as lablab. The publicly available genomic tools and field evaluation data from this study will offer a valuable resource for plant breeders and researchers, to initiate genomic-assisted breeding in lablab which will fast-track genetic gain per unit time and ultimately contribute towards achieving food/nutritional security in the region.

## Introduction

Globally, livestock production provides a prominent source of protein and income/livelihood, particularly among smallholder farmers of Sub-Saharan Africa (SSA) [1, 2]. In SSA, livestock production and productivity are severely restricted by the scarce and low nutritional quality of feedstuffs resulting in poor animal performance and leading to poverty and food/nutrition insecurity [3, 4]. Currently in SSA, feed availability, feedstuff production and utilization practices, forage conservation, improved forage variety development schemes, soil management, use of tools to quantify forage nutritional value, and other aspects of livestock systems are inadequate and need to be upgraded [1, 5, 6]. Most small-scale farmers who produce the majority of the meat and milk in the region rely on rangelands, harvested crop residues and industrial byproducts as their main source of livestock feed, resources that are challenged by their periodic availability and partial suitability for achieving maximum yield from the animals [7, 8]. This challenge is now exacerbated by climate change and dwindling common grazing lands since more land is being allotted for food crops and human settlement [4, 9, 10].

Continued research efforts, particularly focused on developing locally adapted underutilized forage crops known to farmers, can play a critical role in boosting animal performance, leading to an ample supply of animal-sourced foods and ultimately ensuring food/nutritional security in the region [11]. Lablab (*Lablab purpureus* (L.) in the family Fabaceae, is one of the multifunctional crops that originated in Africa [12, 19] and is grown in the tropical and sub-tropical regions of Asia and Africa for fodder and food [13-15]. It is principally self-pollinated with a diploid chromosome number, where 2n = 2x = 22 [16]. It is an annual or short-lived perennial multipurpose forage legume which has versatile uses both as food and feed for animals; or as green manure [17-19]. Interestingly, during the covid-19 pandemic, it was noted that lablab extracts conferred health benefits against viruses such as influenza [20]. Despite all of these applications, limited research attention has been given to lablab to date with no genetic improvement plan recorded.

Lablab was identified, a decade ago, by the African Orphan Crops Consortium (AOCC) as one of the 101 under-researched but highly nutritious African food/feed crops, important for supporting the dietary requirements of consumers and incomes of small-holder farmers in Africa, to have its genome sequenced. This work started [21] and a lablab reference genome (0.5 Gb, 11 chromosomes designated Lp (*Lablab purpureus*)) was published very recently [19]. Exploring the crop’s genetic potential by collecting appropriate germplasm and understanding its diversity status, using genomic tools in combination with phenotyping, is a key approach to fast-track breeding efforts and accelerate the crop’s full domestication [19, 22, 23]. Recently, some agronomic studies have shed light on seed maturity, yield, feed quality traits and intercropping capabilities of lablab [24-27] but there have been few studies on genetic variability and population dynamics [28-31]. For fast-tracking improvement plans, integrated approaches such as phenotyping, genotyping and phenotype-genotype association studies should be carried out and developing genomic tools is an integral part of this endeavor [32-34]. The availability of a high-quality lablab reference genome [19] now enables discovery of genome-wide SNP/Indel markers for targeted marker-assisted breeding.

In this study, a diverse set of lablab accessions, conserved at the International Livestock Research Institute (ILRI) forage gene bank in Addis Ababa Ethiopia, was phenotypically evaluated for morphological and agronomic traits across three locations representing two agroecological settings (low and mid altitudes) in Ethiopia, for two consecutive years and genotyped using a whole genome sequencing approach. The main aim was to: (1) assess genome-wide diversity through phenotypic evaluations and a whole genome sequencing approach, (2) conduct a genome-wide association study and identify SNPs/*k*-mers linked with important agronomic traits and (3) select high yielding and stable genotypes, across the environments, in order to initiate a new breeding program targeted at increasing yield and feed quality traits of this crop.

## Materials and Methods

### Description of the Field Experiment Location and Planting Materials

In this study, a panel of 143 lablab accessions was assembled from collections conserved in the ILRI forage genebank and used for field phenotyping and whole genome sequencing (Supp. Table 1). The selection of these accessions, from a total collection of 340 accessions, was based on passport data and availability of seeds for phenotyping. These materials originated from eight different countries with most accessions donated to the genebank from Kenya and Australia. Seven accessions (6528, 6529, 6533, 6534, 6535, 6536 and 7278) in this panel were of unknown origin and were intentionally included to predict their origin based on their genotypic and phenotypic profile.

The field phenotyping was conducted at three locations in different research stations in Ethiopia, namely Bishoftu geographically located at 8°47’ N, 38°59’ E, and at an altitude of 1890 m.a.s.l., with an Alfisol soil type, average annual rainfall (875mm), temperature (°C) maximum (25), average (19), and minimum (11); Melkassa (located at 8°24’ N, 39°21’ E and at an altitude of 1550 m.a.s.l., with an Alfisol soil type, annual rainfall (875mm), temperature (°C) maximum (23), average (21), and minimum (15)); and Mieso (9°13’ N, 40°44’ E, at an altitude of 1394 m.a.s.l., Alfisol soil type, annual rainfall (875mm), temperature (^0^C) maximum (21), average (18), and minimum (14) (Supp. Fig. 1). For two consecutive years (2020-2021), the trial was conducted under rainfed conditions during the months of July to November.

The field experiment was laid out using a simple lattice design (13*13) with two replications in each location and conducted over two consecutive years. To reduce the effect of soil heterogeneity, twenty-six accessions were duplicated within each replication as controls. Some of these controls are released varieties from different countries. The total number of plots in each replication was 169, partitioned in to 13 incomplete blocks. The spacing between plants and rows was 50 cm and 75 cm, respectively. Land preparation, planting, weeding and other management practices were uniformly applied to all plots across locations. Harvesting of total above-ground biomass was carried out at different times, when 50% flowering had been achieved for each accession.

### Phenotypic Data Collection

After 50 % flowering, eight plants from the central row of each plot were used to record growth and forage biomass yield trait data. Plant height (PH, cm) was measured from the ground base to the tip of randomly selected plants. Fresh leaf weight plant^-1^ (FLW, g), and fresh stem weight plant^-1^ (FSW, g) were measured after separating leaf and stem material. Total fresh weight plant^-1^ (TFWP, g) was determined as the sum of FLW and FSW. The dry leaf weight plant^-1^ (DLW, g), dry stem weight plant^-1^ (DSW, g), and the total dry weight per plant (TDW, g) were measured after oven-drying fresh material at 65 °C for 48 hrs to a constant weight. Number of days to flowering (NoDF) was recorded when an inflorescence was observed in 50% of individuals per plot/accession. The numbers of seeds per pod (NSP) were also recorded, based on those plants for which pods were developing prior to the harvest point being reached. Accession 24778 failed to germinate enough individuals in all locations and both years, and hence it was excluded from phenotypic data analysis.

### Phenotypic Data Analysis

The data were checked for normality using the Shapiro test [35]. The average values for all traits measured for each accession were calculated and normalized ahead of variance comparison (Supp. Fig 2) using the with bestNormalize R package [36]. R statistical software [37] was used to assess the genetic variability. The model used was:

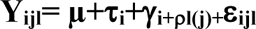

where τ_i_= treatment effect, ψ_i_= replicate effect, πl(j) block within replicate effect, and ijl= random error. A significance level of 5% was used for ANOVA and multiple comparison tests. Among the multivariate methods, the additive main effects and multiplicative interaction (AMMI) analysis was used to account for genotype by environment (GEI) interactions using the metan R package [38]. Measures of mean variance, phenotypic variance, and genotypic variance and coefficients of variation were used to estimate variance components. Phenotypic and genotypic variance, and phenotypic and genotypic coefficients of variation, were estimated following the procedure described previously [39]. For cluster and principal coordinate analysis, the optimum cluster number and membership of accessions within lablab clusters was carried out using the FactomineR R-package and, for visualization, the Factoextra R package was used [40].

### DNA Isolation and Sequencing

Leaf tissues from one-month-old seedlings of the lablab panel accessions were collected (Supp. Table 1) and genomic DNA isolated, using the procedure described by the Qiagen DNeasy® Plant Mini kit (250) (Qiagen Inc., Valencia, CA). Before library preparation, DNA quality was checked on 1% agarose gels and DNA purity was checked using a Nanophotometer® spectrophotometer (IMPLEN, CA, The USA). DNA concentration was measured using a Qubit® DNA Assay Kit in a Qubit® 2.0 Fluorometer (Life Technologies, CA, USA). High-quality DNA with a minimum concentration of 50 ng/µl was used for whole genome sequencing (WGS). All the phenotypically evaluated accessions were sequenced by Novogene, using an Illumina high-throughput sequencer and a paired-end sequencing strategy after library construction.

### DNA Variant Calling and Annotation

Raw reads were trimmed and filtered using the trimmomatic tool [41] to remove low quality and adaptor sequences ahead of mapping to the reference genome. Cleaned reads were mapped to the lablab reference genome [19] using the Burrows Wheller Aligner (BWA) [42]. Samtools was used to convert the aligned reads in SAM format into BAM format [43] followed by filtering out read duplications using Picard tools (http://broadinstitute.github.io/picard). Variant calling against the reference genome was carried out using the Genome Analysis Toolkit (GATK4 v.4.1) [44]. A merged vcf file, containing variants for all the genotyped accessions, was filtered using BCftools (v 1.8) [45]. The SNP quality filtering retained SNPs that are biallelic, polymorphic, with a read depth above 10 and below 300 and according to mapping quality (GQ>20) and minor allele frequency (MAF>=0.05), and a missing rate below 1%. The variant effect predictor program, SnpEff (v 4.3) [46] was used to assess SNPs in coding sequence which may affect the gene product.

### Sequence Data analysis

#### Lablab Genetic Diversity Estimation

Filtered genome-wide SNPs were used for population structure analysis (maximum likelihood estimation of individual ancestries) using the ADMIXTURE tool (v1.3.0) [47]. ADMIXTURE output results were systematically collated using the R pophelper program (v2.3.1) [48], which permits determination of similarities and differences in the ancestral make-up of each population. ADMIXTURE uses a cross-validation procedure by breaking the samples for each *K* value (the number of subpopulations that make up the total population) into five equally sized portions to train the model and estimate the allele frequencies and ultimately predict ancestry assignments of accessions into most probable original populations. Ten-fold cross-validation ’C’ scores were computed for in, ranging from 2 to 10, separately to determine the best fit model for ancestry estimation. A good value of *K* will exhibit a low cross-validation error compared to other *K* values. Furthermore, genomic principal component analysis (PCA) was performed to examine the inter-population distribution using the SNPRelate R package (v1.29.0) [49] and the output was plotted using the Plotly R software package [50]. Finally, phylogenetic trees were constructed using both identity by descent (IBD) and Maximum Likelihood Estimation (MLE) methods and visualized using iTOL [51].

#### Genome-Wide Association Study (GWAS)

For the Genome-Wide Association Study (GWAS), following the exclusion of those that failed to generate enough reads from whole genome sequencing and potential duplicates identified during the diversity analysis, 93 accessions were selected from the panel. Most diverse accessions were included in GWAS analysis in order to minimize false discovery rate (FDR) due to “duplicate” accessions. Best Linear Unbiased Estimator (BLUE) values for each of the measured traits were used for marker-trait association analysis. A marker-trait association was performed for each trait separately with multi-locus GWAS algorithms, Fixed and random model Circulating Probability Unification (FarmCPU) [52], and Bayesian-information and Linkage-disequilibrium Iteratively Nested Keyway (BLINK) model [53] and the Multiple-Locus Mixed Linear Model (MLMM) [54] were implemented in the Genomic Association and Prediction Integrated Tool (GAPIT) version 3 software package within the R environment [55]. The population structure was accounted for by including 3-5 principal components in the subsequent data analysis. The distribution of observed vs. expected −log10(p) values was visualized using Quantile–Quantile (Q–Q) plots to test the fitness of GWAS models for agronomic traits [54]. A threshold *P*-value of 0.001 (–log10(p) > 5) was used to declare significant SNPs for GWAS results. *k*-mer GWAS (kGWAS) analysis was carried out following a protocol described previously [55]. In brief, a 51-bp *k*-mer matrix from Illumina paired raw reads of 136 lablab accessions was created using Jellyfish (v. 2.1.4). Only *k*-mers present in at least two accessions were maintained. Similarly, *k*-mers present in all but one accession were discarded. Each row in the *k*-mer matrix represented the sequence of a 51-bp unique *k*-mer from the entire collection in the first column followed by its presence (1) or absence (0) in each of the accessions in the remaining columns. A “header” file was created following the same order of the accessions in the matrix file. A tab separated file was created with 10^5^ randomly selected *k*-mers extracted from the matrix to account for population structure. kGWAS was run independently for each of the traits as described by others [57]. Associated *k*-mers were mapped to the lablab reference [19] to establish their genomic coordinates. kGWAS methods were sourced from https://github.com/wheatgenetics/owwc/tree/master/kGWAS and running scripts with descriptions from https://github.com/quirozcj/kmerGWAS_descriptions.

## Results

### Field evaluation of a diverse set of lablab accessions across locations and years

The combined analysis of results from the field evaluations indicated significant differences, between lablab accessions and locations, for all traits within and between years. Year was significantly different for leaf and stem fresh and dry weights (Table 1). In addition, the interaction between accession, location and year was significant for all traits indicating that the performance of genotypes was differentially affected by location and year. The mean performance for all traits per accession is presented in Supp. Table 1. Accessions such as 18617, 21043 and 11631 flowered earliest while accessions 21045, 21083 and 24800 were among the late flowering group. The top biomass yielding accessions in terms of TDW were 6536, 14423 and 14439.

**Table 1.** ANOVA summary from combined analysis of eight growth and forage biomass yield traits of lablab accessions evaluated across three locations in Ethiopia.

| Source of variation | NoDF | PH | FLW | FSW | TFWP | DLW | DSW | TDW |
| --- | --- | --- | --- | --- | --- | --- | --- | --- |
| Replication | 17.67ns | 232.9ns | 185662ns | 173230ns | 717791ns | 2066.43ns | 2169.9ns | 8422.5ns |
| Accession | 808.1*** | 3862.6*** | 34853*** | 168718*** | 325233*** | 1597.3 | 7918*** | 14487*** |
| Sites | 27806.5*** | 17210.9*** | 211653*** | 2048028*** | 3464989*** | 3703.4*** | 34647*** | 50376*** |
| Year | 45356ns | 122296 | 2466608*** | 13587224*** | 27713179*** | 120752*** | 537936*** | 1181882*** |
| Accession*site | 190*** | 1197*** | 1197ns | 62217*** | 92667* | 676ns | 3714** | 3070ns |
| Accession*site*year | 103*** | 1113** | 18707*** | 49483ns | 104575* | 806** | 5342*** | 3580* |
| Error | 107.9 | 1062.6 | 14861 | 56144 | 100942 | 737.7 | 3932 | 5965 |

NoDF (number of days to 50 % flowering), PH (plant height), FLW (fresh leaf weight), FSW (fresh stem weight), TFWP (total fresh weight per plant), DLW (dry leaf weight), DSW (dry stem weight), and TDW (total dry weight. Numerical values indicate mean squares, ns (non-significant) \* $p < 0.05$ (5%), \*\* $p < 0.01$ (1%), and \*\*\* $p < 0.01$ (0.1%).

In this study, medium to high genetic variability estimation parameters (phenotypic coefficient of variation (PCV) and genotypic coefficient of variation (GCV)) were recorded for all traits, indicating significant genetic variability (Table 2). Generally, the PCV value was higher than the corresponding GCV value for all measured traits. For example, the PCV value was almost two-fold compared to the corresponding GCV value for DLW, DSW and TDW, indicating that the contribution by the environment was also significant in the expression of these traits. PCV values ranged from 19% to 95% for NoDF and DSW respectively, whereas GCV varied from 15.7% (NoDF) to 48.3% (FSW). A moderate PCV values (19%) was observed for NoDF, whereas high PCV values (>30%) were recorded for the rest of the traits (Table 2). NoDF scored the highest broad-sense heritability (68%) while DSW the lowest (25%). Furthermore, the analysis of expected genetic advance as a percentage of the mean (GAM) indicated that FSW could be improved by 62.9% whilst only a 26.8% alteration could be made for NoDF. TDW could also be improved by 51.5% whilst progress of 41.3 and 49.5% could be made in DLW and DSW, respectively.

**Table 2.** Genetic estimates of lablab accessions based on forage biomass traits evaluated at three different locations of Ethiopia over two consecutive growing years 2021-2022.

| Traits | min | max | mean | $\sigma_g$ | $\sigma_p$ | GCV% | PCV% | H (b) | GA | GAM |
| --- | --- | --- | --- | --- | --- | --- | --- | --- | --- | --- |
| NoDF | 52 | 194 | 97.2 | 233.4 | 341.3 | 15.7 | 19.0 | 0.68 | 26 | 26.8 |
| PH | 55 | 235 | 141.4 | 933.3 | 1995.9 | 21.4 | 31.2 | 0.47 | 43 | 30.1 |
| FLW | 11 | 467 | 188.6 | 6663.9 | 21525.5 | 41 | 73.8 | 0.31 | 93.6 | 47.1 |
| FSW | 20 | 1043 | 391.5 | 37524.5 | 93668.9 | 48.3 | 76.2 | 0.40 | 252.6 | 62.9 |
| TFWP | 32 | 1478 | 583.2 | 74763.7 | 175706 | 45.5 | 69.8 | 0.43 | 367.4 | 61.2 |
| DLW | 5 | 105 | 43. | 286.5 | 1024.3 | 37.9 | 71.8 | 0.28 | 18.4 | 41.3 |
| DSW | 6 | 186 | 72.8 | 1328.6 | 5260.8 | 47.9 | 95.2 | 0.25 | 37.7 | 49.5 |
| TDW | 11 | 292 | 117.6 | 2840.5 | 8805.6 | 44 | 77.5 | 0.32 | 62.4 | 51.5 |

$\sigma_g$ (genotypic variance), $\sigma_p$ (phenotypic variance), GCV (genotypic coefficient of variation), PCV (phenotypic coefficient of variation), H (b) (heritability in a broad sense), GA (genetic advance), GAM (genetic advance as a per cent). NoDF (number of days to 50 % flowering), PH (plant height), FLW (fresh leaf weight), FSW (fresh stem weight), TFWP (total fresh weight per plant), DLW (dry leaf weight), DSW (dry stem weight), and TDW (total dry weight).

### Principal Component and Cluster Analysis for forage biomass based agro-morphological traits

Principal component analysis (PCA) and cluster analysis were performed to produce a cluster that shows the approximate relationships among a set of accessions and to convert a set of observed traits of possibly correlated variables into a set of values of linearly uncorrelated standardized quantitative variables. In this analysis, divergence was observed based on growth and forage biomass traits which identified the grouping through the measured traits response.

Principal component analysis (PCA) identified two principal components with Eigen values of greater than one which contributed 83.13% of the cumulative explained variation from the first two components (Supp. Table 2). The first principal component (PC1) accounted for 66.78% of the total explained variance, with a significance loading of NoDF (0.33), PH (0.46), FLW (0.86), FSW (0.95), TFWP (0.97), DLW (0.89), DSW (0.85), TDW (0.96) and all the traits were positively correlated with each other. Similarly, the second principal component (PC2) accounted for 16.34% of the total explained variance. A PCA biplot shows the degree of correlation among measured traits and a tight angle between vectors is an indicator of a high and positive correlation (Fig. 1A). The strongest correlation was found between TDW and FSW. Likewise, DLW and TFWP were highly correlated. NoDF and PH were the furthest from the rest of the measured traits but still had a positive correlation with the rest of the traits. Based on standardized data from two years of mean values of eight quantitative traits, the cluster analysis revealed four basic clusters with cluster I mainly composed of 2-seeded lablab accessions (Fig. 1B). All accessions identified to be high yielding (TDW/TFWP) (Supp. Table 3) were in cluster IV, except for accession 11615.

**Fig. 1.**
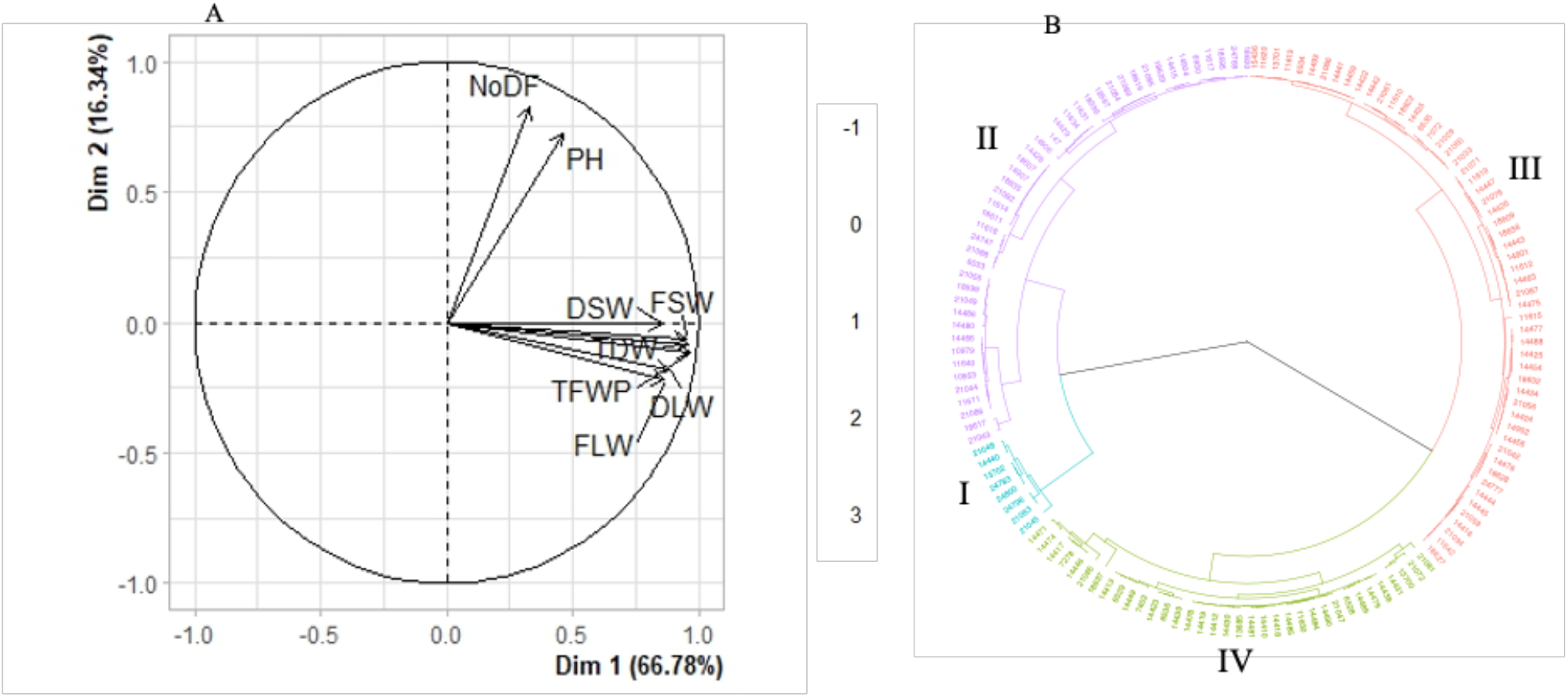
Variable correlation plot showing the relationships among eight quantitative traits in 142 lablab accessions (A) and Cluster Analysis of phenotyped lablab accessions based on forage biomass traits (B)

### Performance, Yield and Stability of lablab Accessions

The AMMI analysis of variance indicated highly significant (*P<0.01*) accession by environment interaction effects for all traits. Genotypic factors accounted for a larger proportion of the treatment mean of squares for PH, NoDF, and TDW. The environment was significant for TFWP, NoDF, DSW, and FSW but not for the other traits (Supp. Table 4).

Additive main effect and multiplicative interaction stability analysis ranked the accessions based on AMMI stability variance index (ASI) score for TDW (Supp. Table 1: AMMI score). Low scores in ASI represent stable accessions for that particular trait across tested environments. Hence, in this study accession 21048 scored the lowest ASI score indicating it was the most stable accession for TDW, whereas accession 18622 was ranked the least stable with the highest ASI score for this trait. The combination of yield and stability rankings (SSI) indicated that accessions such as 14423 have high yield with more or less stable performance across the three locations, in contrast to 6529 which exhibited high mean TDW but showed lower stability across the locations. Therefore, the breeders’ aim of selecting high yielding accessions across multiple environments could be met with priority accessions such as 14423, 14439 and 6536.

### Whole Genome Sequencing of Lablab Accessions

Although all phenotyped lablab accessions were genotyped seven failed to generate enough sequence reads and hence were excluded from the diversity analysis. More than 2 billion reads were generated for the remaining 136 lablab accessions with an average of ca. 15 million reads/accession (Supp. Table 1). The average coverage depth of all the bam files was 7.7. More than 20 million raw variants (SNPs: ca. 16 million; Indels: ca. 3 million) were detected when the trimmed reads were mapped against the lablab reference genome. The SNPs were filtered for downstream analysis based on polymorphism (bi-allelic only), read depth (between 10 and 300), genotyping quality (GQ >20), minor allele frequency (MAF > 0.05) and missing rate (< 1%). Half a million SNPs passed the aforementioned threshold (Supp. Fig. 3) and were retained for measuring genetic diversity parameters and the association mapping study. Functional annotation using SnpEff revealed that 42% of the SNPs fall in the intergenic region of the lablab genome. Furthermore, 53% of the SNPs were missense, 0.7% were nonsense and the rest were silent polymorphisms. In terms of putative impact, ca. 5% of the variants were predicted to have low to high impact and most of the SNPs (95%) fell under the modifier category (Supp. Fig. 4).

Different numbers of SNPs were detected on different chromosomes, with the highest number of SNPs recorded for Lp03 (78,408) and the lowest for Lp08 (36,384) (Supp. Table 5/Supp. Fig. 3). This is not unexpected as Lp03 is the largest chromosome in the reference genome [19]. Interestingly, the highest SNP density rate was recorded for Lp10 and the lowest for Lp07 (Supp. Table 5). On average a SNP was detected at ca. 800 bases for all chromosomes. Chromosomes Lp5, Lp7 and Lp11 scored the highest MAF (0.24) and the lowest was recorded for Lp09 (Supp. Table 5). A similar trend among the eleven lablab chromosomes was also observed for polymorphic information content (PIC, Supp. table 5). A rarefaction curve indicated that nearly 100% of unique SNPs could be picked using approximately half of the panel (e.g. 95% unique SNPs were detected with 60 samples)(Fig. 2).

**Fig. 2.**
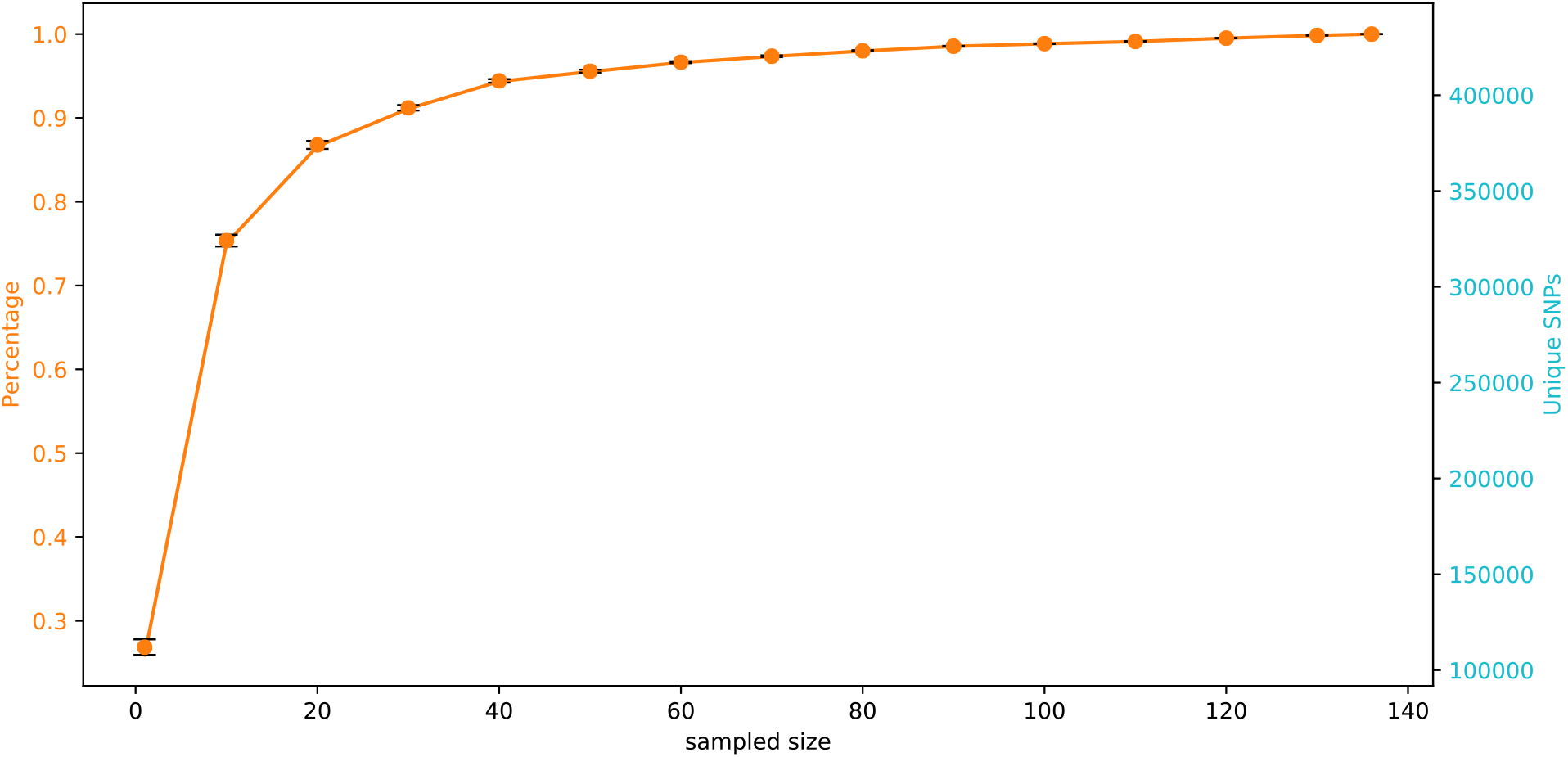
Genic SNPs rarefaction curve per 100 replicates

### Population Structure and Clustering Among Genotyped Lablab Accessions

We utilized a clustering approach based on the maximum likelihood estimation (ADMIXTURE software), using half a million filtered SNPs distributed over the 11 lablab chromosomes, to detect underlying ancestral populations of the 136 lablab accessions. The ADMIXTURE simulation demonstrated that a *K* value of 8 exhibited lowest cross validation error at *K* = 8, inferring that the 136 analyzed accessions belong to eight underlying subpopulations with the highest likelihood (Supp. Fig. 5). This predicted population structure displayed partial membership of more than one population for some accessions whereas others showed membership of only one population. For example, 21086, 21056, 21081 and 24796 grouped in a single cluster using *k* = 8. On the other hand, *K* = 2 with the highest cross validation error clustered the accessions into two clear distinct groups. The PCA analysis showed a similar trend where PC1 separated the panel into two groups. (Fig. 3). A phylogenetic tree generated by hierarchical clustering provided corroborating evidence to support the ADMIXTURE analysis in that there was no clear segregation of accessions based on geographic origin (Supp. Fig. 6).

**Fig 3.**
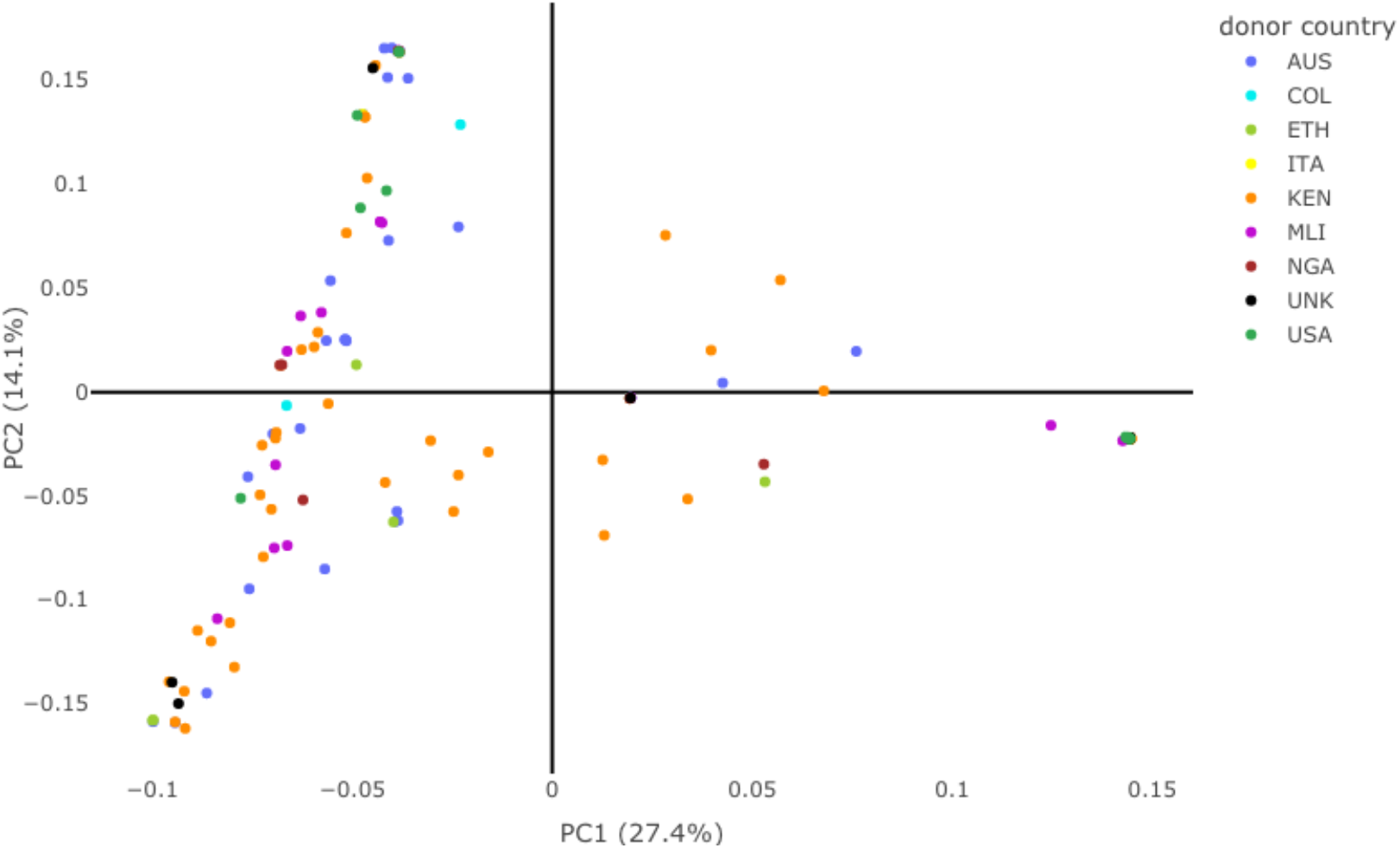
PCA plots of the first two principal components of 136 sequenced lablab accessions. (3d version: file:///Users/Abel/Documents/JIC%20migrated%20docs/A%20FLAIR%20ROAD%20MAP/A% 2075K%20lablab%20project/lablabSNP/lablab.pca/lablab.136pca2.3d.html

Several accessions that were a member of a single cluster in the ADMIXTURE analysis also clustered similarly in the phylogenetic tree. Core hunter analysis with ca. half a million SNPs predicted 27 accessions can represent the diversity among all of the 136 accessions (Supp. Table 1: Core Hunter selection). Core Hunter analysis was further used to predict different numbers of core accessions that best represent the panel in this study (Supp. Table 1). From the seven accessions of unknown origin, three (6529, 6533 and 6535) clustered together in the phylogenetic tree and showed single cluster membership in ADMIXTURE analysis, indicating that they are highly related, but the rest were clustered into different clusters. The accessions 6533 and 6535 also clustered together in the phylogenic tree, based on agro-morphological traits.

### Principal component analysis based on SNP marker set

The PCA revealed an interesting trend with two major clusters and a scattered set of materials, presumably landraces or wild materials but no clear separation of accessions based on region of origin (Fig. 3). The accessions in this panel were obtained from eight donor countries, but the PCA did not reveal ecotyping based on their origin. Nonetheless, each one of 127 accessions has a unique set of SNPs that can be utilized to develop molecular barcodes for precise identification in the future (Supp. Fig. 7).

### Genome Wide Association Study (GWAS)

A GWAS analysis was carried out to identify genetic variants (SNPs and *k*-mers) significantly associated with agro-morphological traits measured in the present study. In the SNP-based GWAS, using 93 lablab accessions, all traits showed significant associations higher than a −log10(p) value of 6 for at least one genomic location (Fig. 4). Likewise, in the *k*-mer based GWAS, using 136 lablab accessions, strongly associated *k*-mers were identified for all traits, except for PH (Supp. Table 1: *k*-mer-GWAS). Interestingly, nearly 200 *k*-mers were significantly associated with DLW (> −log10(p) value of 6) which was a significantly higher number of *k*-mers than recorded for other traits. Some *k*-mers identified were negatively associated with target traits, implying that these markers contribute negatively to the trait; for example, all significantly associated *k*-mers for NoDF were associated with early flowering, i.e., displayed negative correlation with this trait. Interestingly, the only *k*-mer significantly associated with FSW was negatively correlated with this trait (Supp. Table 1: *k*-mer-GWAS).

**Fig 4.**
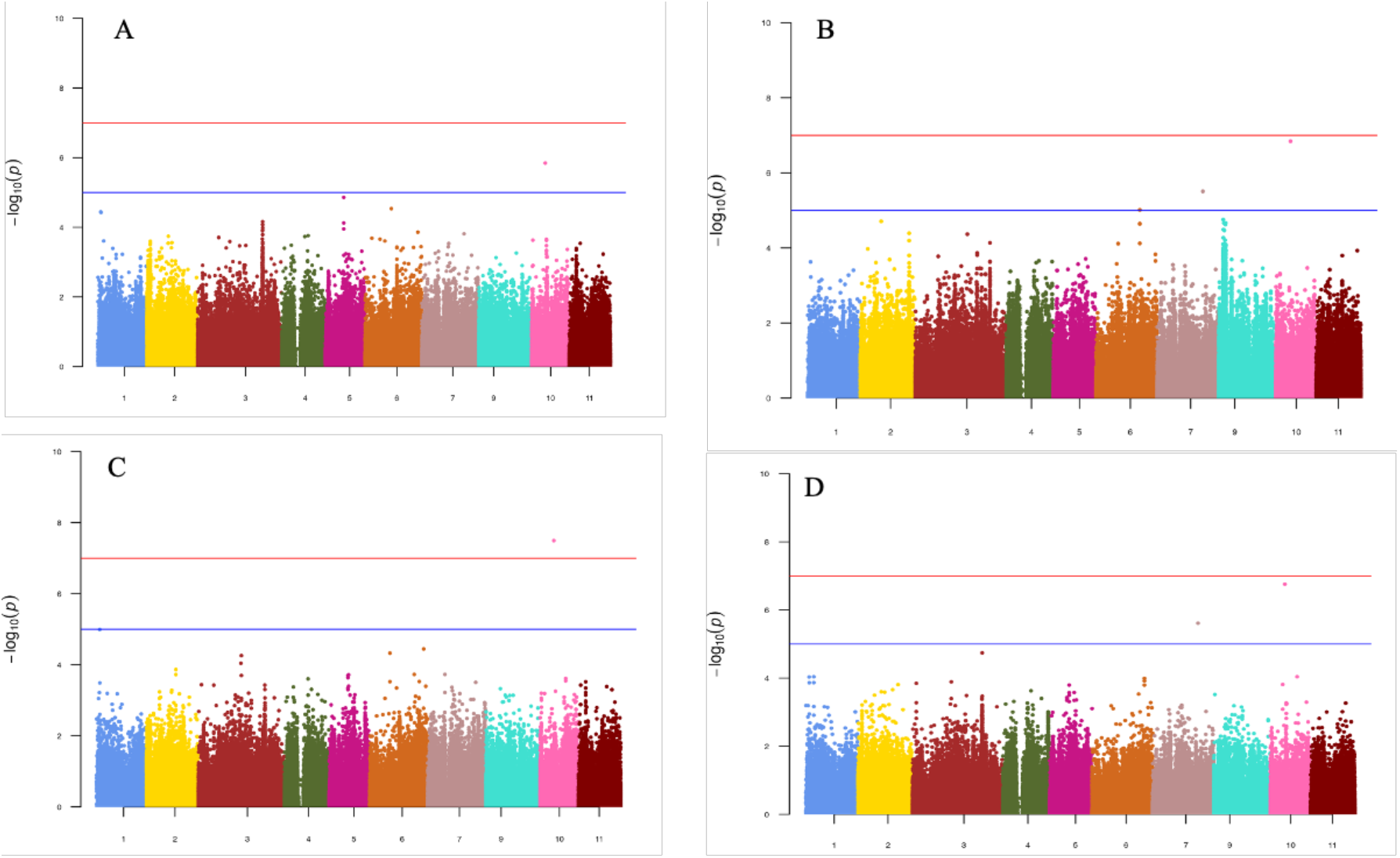
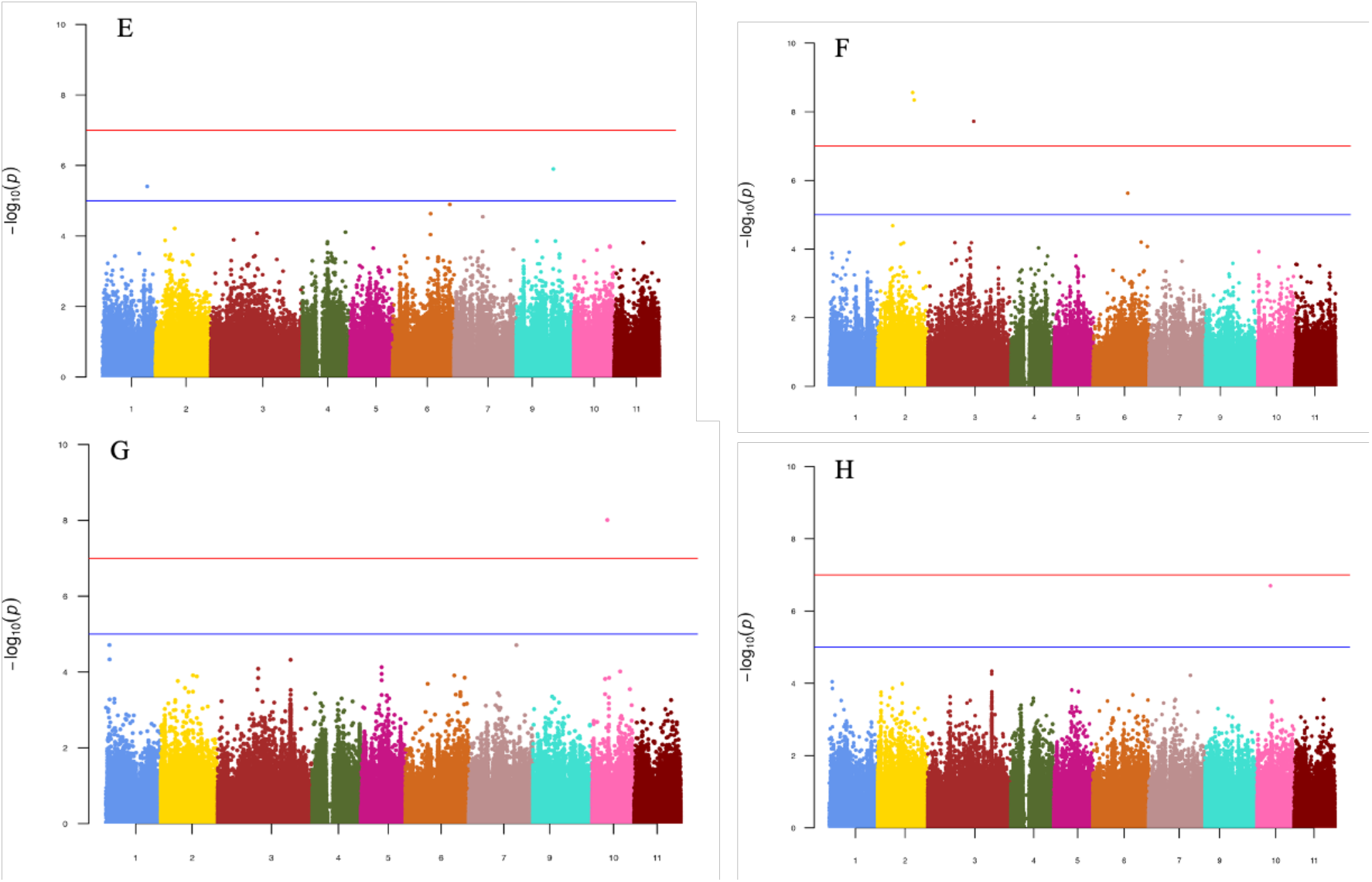
GWAS Manhattan plot SNPs identifying QTLs associated with eight agro-morphological traits evaluated over two years for 93 diverse lablab accessions. (A) DLW (dry leaf weight), (B) DSW (dry stem weight) (C) FLW (fresh leaf weight), (D) FSW (fresh stem weight) (E) NoDF (number of days to 50 % flowering), (F) PH (plant height), (G) TFWP (total fresh weight per plant), and (H) TDW (total dry weight)

## Discussion

In this study, a panel of 142 lablab accessions, conserved by the ILRI genebank, was phenotyped across three different locations in Ethiopia and over two years, revealing different levels of variability and heritability among agronomic performance traits. The same accessions were genotyped using a whole genome sequencing approach that generated high-density genome-wide markers which were used for diversity assessment and marker-trait association studies.

### Field Characterization of Lablab Accessions across Locations and Years

In the present study, significant variation (*p <* 0.01) was observed for all eight measured traits indicating the existence of high genetic diversity among the panel of accessions (Table 1). Variations in response to different environments by individual accessions was observed, similar to that reported previously [15, 58]. The mean performance of all accessions across the three environments also varied significantly (*P <* 0.05), highlighting the environmental influence on the expression of those traits (Table 1). The phenotypic traits studied here delineated the accessions into four main sub-clusters with a variable number of members/accessions within each cluster (Fig 1). Interestingly, this clustering was able to grouped all 2-seeded accessions (13702, 14440, 21045, 21083, 24796 and 24800) into cluster I along with two 4-seeded accessions (24783and 21048), a similar finding to that reported previously [19, 58, 59]. Furthermore, clustering based on morphological traits placed most of the best-performing accessions, in terms of TDW, except 11615, into cluster IV (Fig 1) which makes this cluster a preferred source of choice parental lines for a new cycle of breeding programmes targeting improvement in forage performance. However, in future hetrosis breeding programs accessions such as 11615 could also play a key role in improving yield traits while minimizing inbreeding depression due to its divergent genetic background.

Genotypic variability is a precondition for the advancement of any breeding program and among variability parameters genotypic and phenotypic variance estimates are good indicators of how much improvement can be achieved during genetic selection [60]. PCV and GCV values are classified as low (when < 10%), medium (between 10% and 20%), and high (>20%), according to Deshmukh et al. [61]. High PCV and GCV values i.e. > 20% were observed for all traits except for NoDF, a similar result to that reported previously [60] (Table 2). Furthermore, the difference between PCV and their respective GCV values were relatively small for NoDF and PH, indicating that these traits were less influenced by the environment and potentially more amenable to improvement through phenotypic selection. Early flowering is a key trait, particularly in lowland areas where rainfall periods are short. A number of plants, including legume species, are known to deploy drought escape mechanisms, i.e., completing their life cycle quickly in order to avoid water stress during the later growth stages [63, 64]; hence early maturing accessions identified in this study could be useful candidates for areas with short rainy periods. For example, Meiso is located at a relatively lower altitude (1550 m.a.s.l.) and known for its short rainfall span; therefore, accessions such as 18600, 14429 and 24768 with medium NoDF and average forage yield can be selected for such localities.

NoDF (68%) and PH (47%) showed the highest broad-sense heritability in the present study which implies that the performance of these traits was principally influenced by additive gene interactions and hence selection will be efficient for improvement of these traits. However, broad sense heritability (H) includes both additive and non-additive components of the variation and hence does not always provide a full indication of genetic gains that can be made through selection [65, 66]. Therefore, it is necessary to estimate the genetic advance as a percentage of the mean (GAM) to determine the actual progress. In this study, NoDF and PH scored 26.8% and 30.1% (GAM) respectively which indicated the progress that can be made through selection of early or late maturing accessions in the next generation.

The additive main effects and multiplicative interaction (AMMI) models are the most suitable in these studies when interaction between the environment and genotypes is significant such as in the present study [67]. Stable genotypes normally have low AMMI stability index (ASI) scores while unstable ones have high ASI scores [68]. However, stability in performance for yield or other traits does not necessarily warrant selection as in some scenarios the most stable genotypes do not always have the best performance and vice versa [69, 70]. AMMI analysis identified accessions 14423 and 14440 as two of the highest yielding accessions in terms of TDW with relatively low ASI scores, which implies that more of an additive genetic effect is the major driving factor (Supp. Table 1. AMMI score). Subsequently, we have selected high-ranking accessions, based on TDW/TFWP (Supp. Table 3) and these accessions are currently undergoing seed multiplication for region-wide tests across East Africa, leading to the possibly of variety releases in the coming few years. Accession 6536, a wild accession, scored the second highest TDW and relatively good stability score. This accession and a few others performed as well in terms of total dry weight (TDW), compared to released varieties (Supp. Table 3).

### Genomic Tools for Lablab Improvement

The reduction in the cost of Next-Generation Sequencing (NGS) technology has greatly facilitated the deciphering of complex genomes. Unfortunately, such genomic tools are scarce among key tropical forage species such as lablab. In the current study, a global panel of 136 lablab accessions, sourced from eight different countries, was genotyped at the whole-genome level and, using the recently released reference genome, millions of SNPs were detected. On average a SNP was identified every 800 bases for all chromosomes (Supp. Table 5) which is a much higher SNP density than that reported for lablab recently (one GBS-based SNP markers per ca. 100Kb) [19, 28]. Despite the high number of genome-wide SNP markers identified in the present study, the rate of unique SNPs identified decays with half of the genotyped accessions (Fig 2) which implies possible duplications in our panel. Interestingly, Core-Hunter analysis predicted that 27 accessions (ca. 20% of the total analysed) can represent the overall SNP diversity in the panel (Fig 2). A previous core collection analysis in lablab revealed a similar trend where only 47 accessions (ca. 10%) represented a panel of nearly 500 lablab accessions [12].

### Genetic Diversity among Lablab Accessions

To clarify the phylogenetic relationship and population genetic structure of cultivated and wild lablab accessions, ADMIXTURE analysis, PCA and hierarchical clustering approaches were employed and none of these approaches clustered accessions according to their country of origin, implying exchange of materials between countries or genebanks. A number of diversity studies on lablab based on agro-morphological traits and molecular markers reported similar findings [26, 71, 72]. Genetic assignment analysis using ADMIXTURE predicted eight sub-populations (optimal value of *K* = 8) with a high level of population admixture between genetic groups (Supp. Fig 5). Most accessions that showed single cluster membership in ADMIXTURE analysis also clustered together in a hierarchical cluster analysis (Supp. Fig. 6). The genomic tools developed in this study predicted possible genetic groups for the seven accessions of unknown origin which provides valuable information for the future genebank management of these accessions. Furthermore, these genomic tools have facilitated the identification of unique SNP/s for 93% of genotyped accessions which can be used as molecular barcodes in future breeding programs and in genebank management (Supp. Fig. 7).

A previous genetic diversity study on lablab, using a genotyping-by-sequencing approach, which included a significant number of the accessions in this panel predicted two main sub-populations composed of mainly 2-seeded and 4-seeded accessions, respectively [19]. In the present study, clustering based on SNP data predicted two major clades, where one cluster was composed of only five accessions of which two were 2-seeded. In the SNP based clustering, two 2-seeded lablab accessions (14440, 21045), ssp. *uncinatus*, uncharacteristically clustered with 4-seeded accessions which is not in line with what has been reported previously [19, 72] and hence needs further investigation further. However, phylogenetic analysis based on agro-morphological data clustered all 2-seeded accessions into cluster I (Fig 1) with only two 4-seeded accessions (24783 and 21048) included as predicted previously by Njaci et al. [19]. Interestingly 24783, is a 4-seeded lablab accession which clustered with the 2-seeded accessions in both SNP and morphological data based phylogenetic trees (Fig 1) while 21048 was not successfully genotyped. This accession, 21048, is classified as ssp. *uncinatus* which could partly explains its segregation from the rest of 4-seeded accessions. Kongjaimun et al. [12] reported that some of the 4-seeded lablab accessions are closely related to wild 2-seeded accessions and this could be a similar result. Both hierarchical and PCA clustering, based on a SNP marker dataset, revealed high similarity among the panel, though sourced from different genebanks which implies possible duplication in the panel.

### Novel Genomic Resources to Identify Candidate Loci in Lablab

Genome wide association studies (GWAS) are common in the NGS era in order to identify linkage disequilibrium between phenotypic traits and genetic markers (SNPs/Indels) without any prior knowledge of population structure or kinship matrix of a panel [73]. Previously, a GWAS analysis in lablab has identified QTLs for plant height (PH) in chromosomes Lp3 and Lp4 [19]; the present study has identified additional QTLs on the same chromosomes and further on chromosomes 6, 7 and 8 (Fig 4). PH is one of the key traits determining forage quality, yield, and management strategy (harvesting and/or grazing) [74, 75]. In the field evaluations, PH showed the highest broad-sense heritability (Table 2) which implies potential for improvement and the QTL identified for PH in this study can play a significant role in future selection efforts. Floral induction is triggered by environmental cues such as day length and temperature and genetic factors associated with this trait are key to determining biomass yield, feed quality and response to water stress [76-78]. In this study, two QTL, on chromosomes Lp1 and Lp9, were significantly associated with NoDF (Fig 4). However, Njaci et al. [19] reported QTL on a different chromosome for the same trait suggesting that different regions of the genome may be impacting on this trait.

When whole genome sequence data are available, such as in this study, *k*-mer-based GWAS allows identification of variation that cannot be covered with commonly used SNP-based GWAS [79-81]. *k*-mer-based analyses allow the detection of *k*-mers originating from regions that are absent from the reference, such as large insertions, and also help minimize false positive associations arising from errors during variant calling and/or genotyping steps [80]. In this study, *k*-mers significantly associated with all target traits except for PH were detected (Supp. Table 1: kmer-GWAS) and to our knowledge this is the first report of a *k*-mer based association study for a tropical forage crop. In summary, valuable associations were identified for most key traits, with both *k-*mer and SNP based GWAS, which will enable fast-track selection of these traits in future breeding programs.

## Conclusion

Lablab is one of the key tropical forages grown in the region and, in this study, we have presented a comprehensive analysis of a diverse panel of lablab accessions grown under two environmental conditions leading to the identification of best-performing accessions with high fresh and dry forage biomass production, under low to medium precipitation environments. Furthermore, the genomic tools we have developed provide an unprecedented public resource for lablab genomic research and offers the opportunity to reduce the duration of the breeding cycles in plant improvement programs through the application of marker-assisted selection (MAS) and/or genomic selection (GS) and mutagenesis studies of trait enhancement.

## Supporting information

Metadata

## Acknowledgments

The authors would like to thank Sergei Kushnir and Noel Ellis for reading and commenting on the original manuscript. We also would like to thank Noel Ellis for the discussion to improve diversity analysis results.

## Author contribution

TA, CD, MA and CJ designed and supervised the project. Funding was acquired by TA, MA, CJ and CD. LH collected leaf samples and extracted DNA. MA, LH, TA, MS and HE involved in phenotype data collection and supervision of the field trials. CJ involved in the supervision of the genotyping. TA, LH, QCJ and CJ participated in phenotypic and genotypic data summary. TA, LH. NE and CJ wrote the manuscript. All authors contributed to manuscript revision, read, and approved the submitted version.

## Declarations

Ethics Approval and Consent to Participate Not applicable.

## Consent for Publication

Not applicable.

---

**Supp. Table 2.**
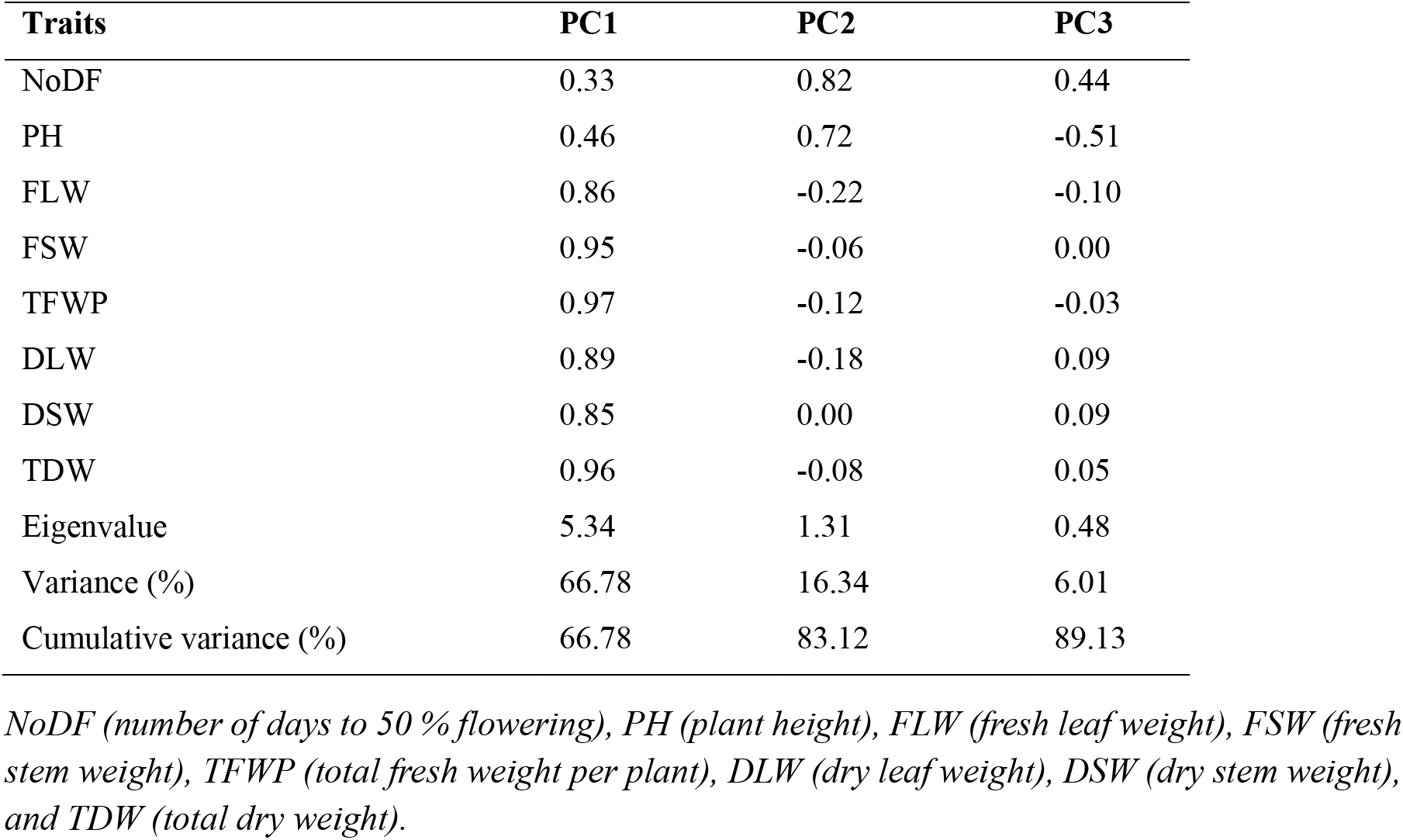
Traits contributing the highest components of variation among lablab accessions.

**Supp. Table 3.**
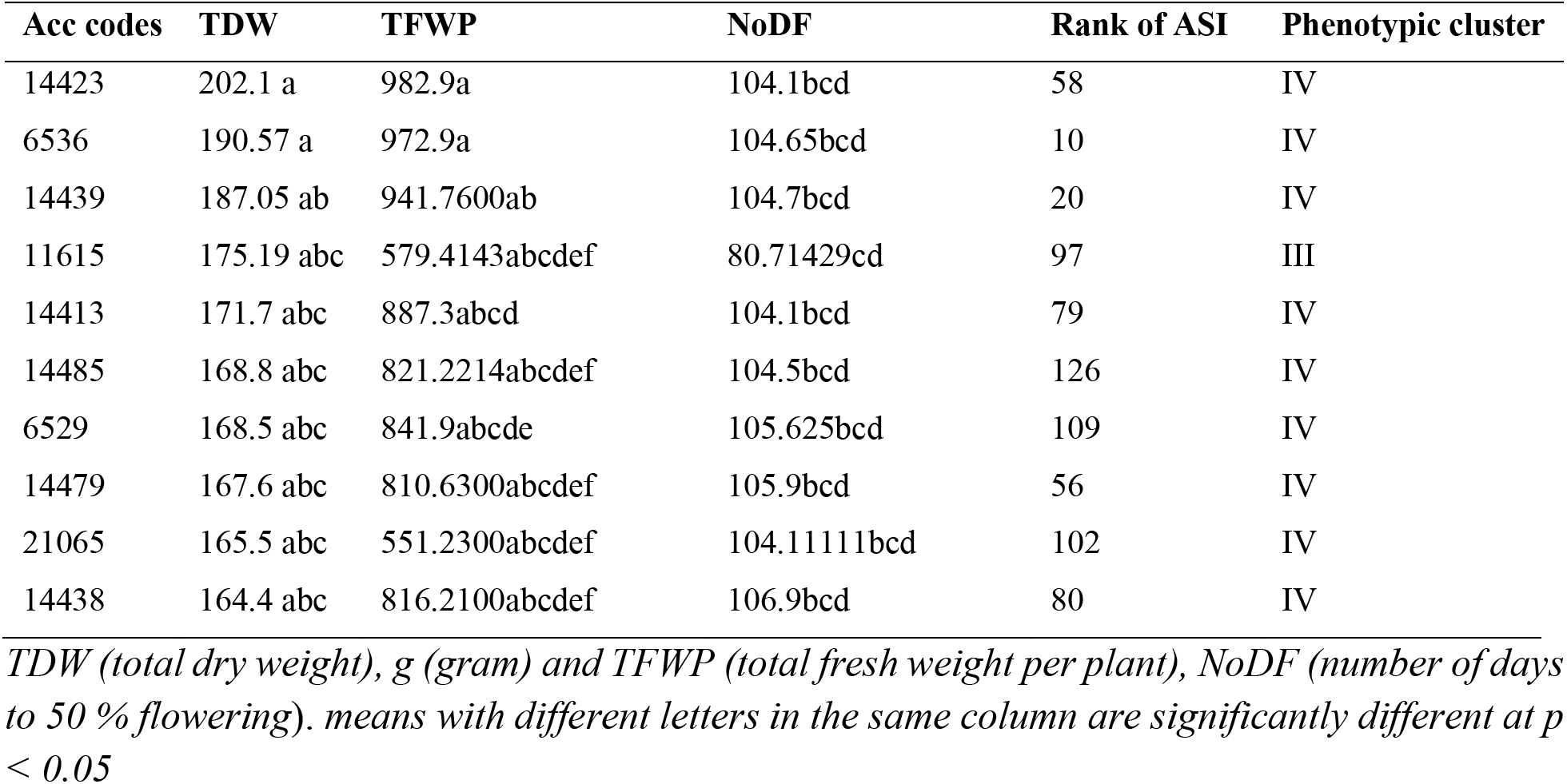
Best-performing accessions based on high yield (TDW & TFWP)

**Supp. Table 1.** AMMI analyses of variance for 142 lablab accessions evaluated for two consecutive years in three growing agro-ecologies of Ethiopia for growth, forage biomass-related traits.

| Traits | Source of variations | Environment | Replication<br>(Environment) | Accession | Accession*Environment | IPCA1 | IPCA2 | Error | Total |
| --- | --- | --- | --- | --- | --- | --- | --- | --- | --- |
| NoDF | Df | 2 | 3 | 142 | 269 | 143 | 141 | 1114 | 1814 |
|  | Sum Sq | 52040.79 | 0.2 | 95265.34 | 313.8 | 253.4 | 91.9 | 11697.22 | 159662.9 |
|  | Mean Sq | 26020.4*** | 0.098ns | 670.8*** | 1.1ns | 1.7 | 0.6 | 10.5002 | 88.01703 |
|  | F value | 265409.5 | 0.09 | 63.89238 | 0.11 | 0.17 | 0.06 |  |  |
| PH | Df | 2 | 3 | 142 | 267 | 143 | 141 | 1097 | 1795 |
|  | Sum Sq | 20205.7 | 6941.8 | 252356.53 | 2457.3 | 1520.1 | 578. | 88749.6 | 372809.2 |
|  | Mean Sq | 10102.85ns | 2313.96*** | 1777.1*** | 9.2ns | 10.6 | 4. | 80.9 | 207.6 |
|  | F value | 4.356 | 28.6 | 21.966764 | 0.11 | 0.13 | 0.05 |  |  |
| FLW | Df | 2 | 3 | 142 | 267 | 143 | 141 | 1084 | 1782 |
|  | Sum Sq | 255408.99 | 76033.42 | 1784581.2 | 57118.06 | 32268.84 | 14076.3 | 1798146.8 | 4017633.6 |
|  | Mean Sq | 127704.4ns | 25344.4*** | 12567.4*** | 213.9ns | 225.6 | 99.8 | 1658.807 | 2254.5 |
|  | F value | 5.0387514 | 15.2 | 7.576212 | 0.12 | 0.14 | 0.06 |  |  |
| FSW | Df | 2 | 3 | 142 | 268 | 143 | 141 | 1102 | 1801 |
|  | Sum Sq | 3924817.79 | 260602.17 | 16298760 | 336144.08 | 212818.1 | 82066.1 | 11524318 | 32639526 |
|  | Mean Sq | 1962408.8** | 86867.3*** | 114779.9*** | 1254.2ns | 1488.2 | 582.0 | 10457.6 | 18123 |
|  | F value | 22.5 | 8.3 | 10.975709 | 0.12 | 0.14 | 0.06 |  |  |
| TFWP | Df | 2 | 3 | 142 | 268 | 143 | 141 | 1097 | 1796 |
|  | Sum Sq | 6922049.3 | 779579.2 | 28221117 | 696615.2 | 434083.3 | 166561 | 23070902 | 60290907 |
|  | Mean Sq | 3461024.6* | 259859.7*** | 198740.2*** | 2599.3ns | 3035.5 | 1181.3 | 21030.9 | 33569.5 |
|  | F value | 13.3 | 12.3 | 9.4499154 | 0.12 | 0.14 | 0.06 |  |  |
| DLW | Df | 2 | 3 | 142 | 268 | 143 | 141 | 1089 | 1788 |
|  | Sum Sq | 3145.6 | 2590.674 | 88656.33 | 3218.068 | 2053.3 | 639.7 | 119184.14 | 219487.8 |
|  | Mean Sq | 1572.79ns | 863.5*** | 624.3*** | 12.0ns | 14.3 | 4.5 | 109.4 | 122.7 |
|  | F value | 1.8 | 7.89 | 5.704674 | 0.1 | 0.13 | 0.04 |  |  |
| DSW | Df | 2 | 3 | 142 | 267 | 143 | 141 | 1097 | 1795 |
|  | Sum Sq | 53973.042 | 5531.4 | 477753.69 | 9909.55 | 6129.8 | 2260.14 | 356402.7 | 911960.47 |
|  | Mean Sq | 26986.52* | 1843.8*** | 3364.4*** | 37.11ns | 42.8 | 16.02 | 324.8886 | 508.05 |
|  | F value | 14.6 | 5.6 | 10.355742 | 0.11 | 0.13 | 0.05 |  |  |
| TDW | Df | 2 | 3 | 142 | 268 | 143 | 141 | 1104 | 1803 |
|  | Sum Sq | 78830.8 | 14741.4 | 1040406.3 | 24625.4 | 15675.13 | 5062.76 | 924380.37 | 2103722.2 |
|  | Mean Sq | 39415.4ns | 4913.81*** | 7326.8*** | 91.8ns | 109.6 | 35.9 | 837.3 | 1166.7899 |
|  | F value | 8.021353 | 5.8 | 8.7505022 | 0.1 | 0.13 | 0.04 |  |  |

**Supp. Table 5.** Total number of filtered SNPs detected for 136 lablab accessions and genetic diversity parameters per chromosome.

| **Chromosome** | **Chromosome**<br>**length (#)** | **SNP base**<br>**(#)** | **SNP Variants**<br>**rate** | **SNP**<br>**Density/chr** | **MAF** | **PIC** |
| --- | --- | --- | --- | --- | --- | --- |
| Lp01 | 36,690,027 | 38,549 | 951.8 | 0.10 | 0.22 | 0.25 |
| Lp02 | 32,207,972 | 37,478 | 859.4 | 0.12 | 0.21 | 0.25 |
| Lp03 | 63,374,807 | 78,408 | 808.3 | 0.12 | 0.22 | 0.25 |
| Lp04 | 32,757,788 | 48,103 | 681.0 | 0.15 | 0.23 | 0.26 |
| Lp05 | 29,843,965 | 37,998 | 785.4 | 0.13 | 0.24 | 0.27 |
| Lp06 | 42,481,378 | 53,850 | 788.9 | 0.13 | 0.21 | 0.24 |
| Lp07 | 42,956,154 | 40,408 | 1063.1 | 0.09 | 0.24 | 0.26 |
| Lp08 | 38,128,793 | 36,384 | 1048.0 | 0.09 | 0.22 | 0.25 |
| Lp09 | 39,552,585 | 51,573 | 766.9 | 0.13 | 0.18 | 0.22 |
| Lp10 | 28,770,402 | 43,351 | 663.7 | 0.15 | 0.22 | 0.25 |
| Lp11 | 31,106,568 | 39,110 | 795.4 | 0.13 | 0.24 | 0.26 |
| **Total** | **417,870,439** | **505,212** |  |  |  |  |

*MAF (minor allele frequency), PIC (polymorphic information content)*

## Supplementary figures

**Supp. Fig 1.**
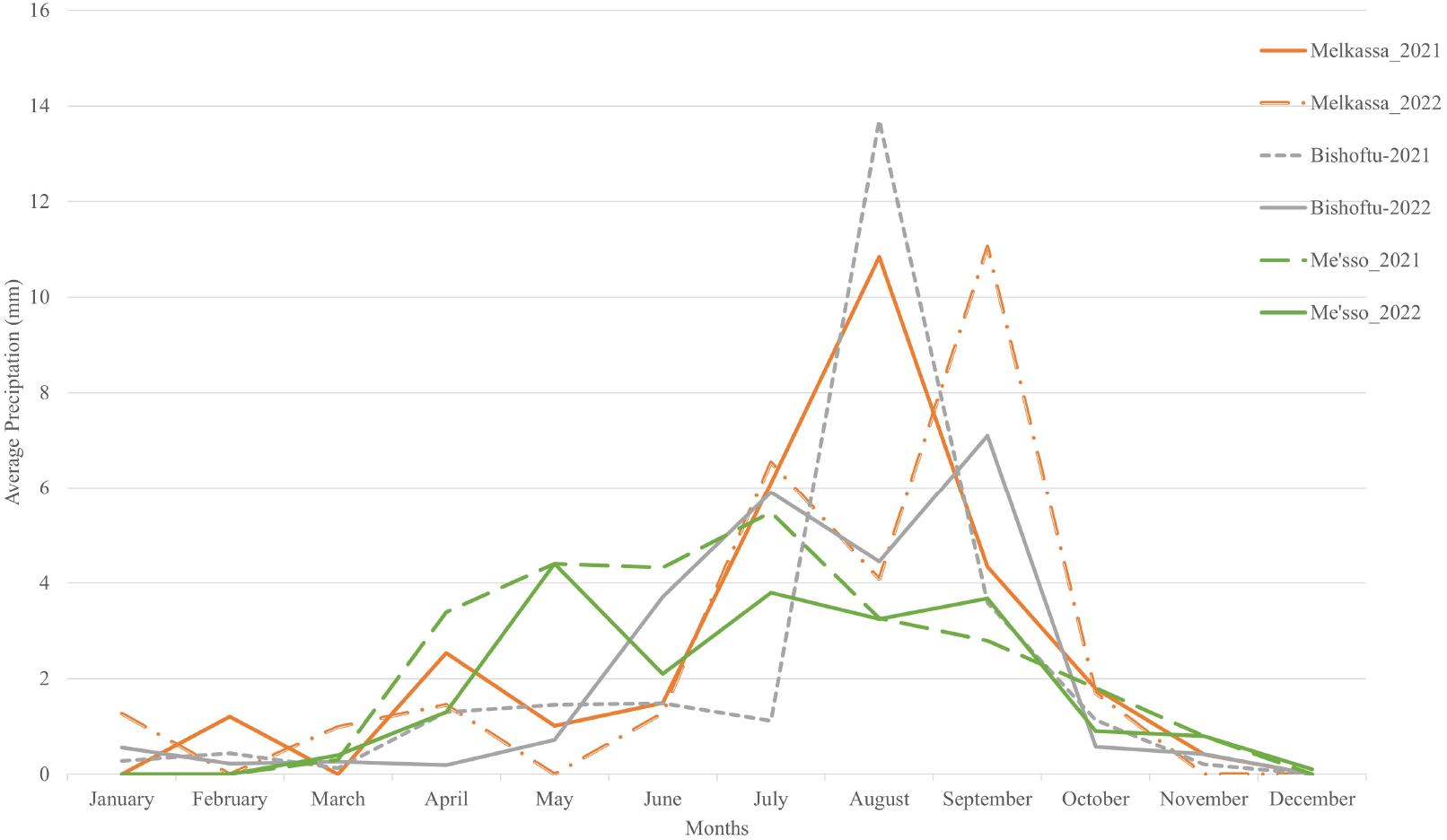
Monthly average precipitation distribution from 2021-2022 for each site

**Supp. Fig 2.**
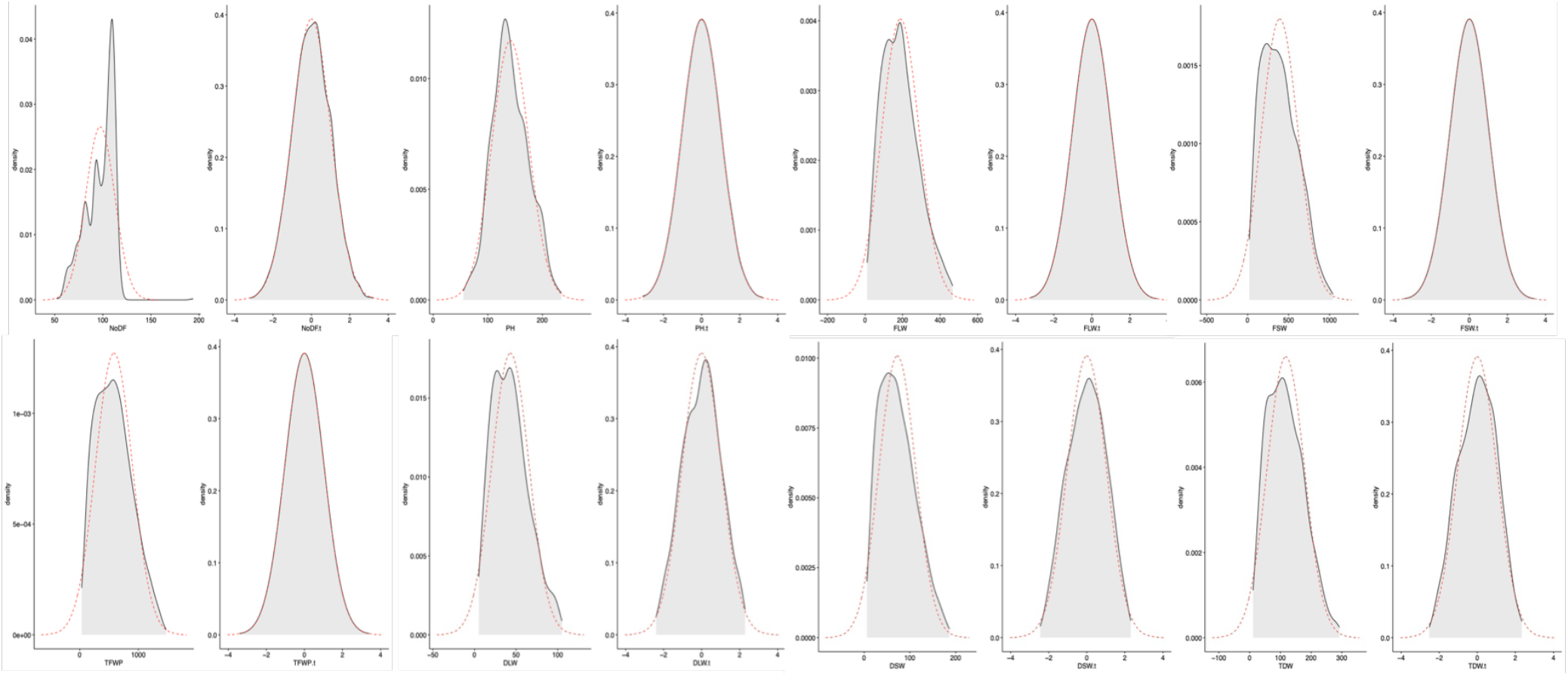
Distribution of traits before and after transformation

**Supp. Fig 3.**
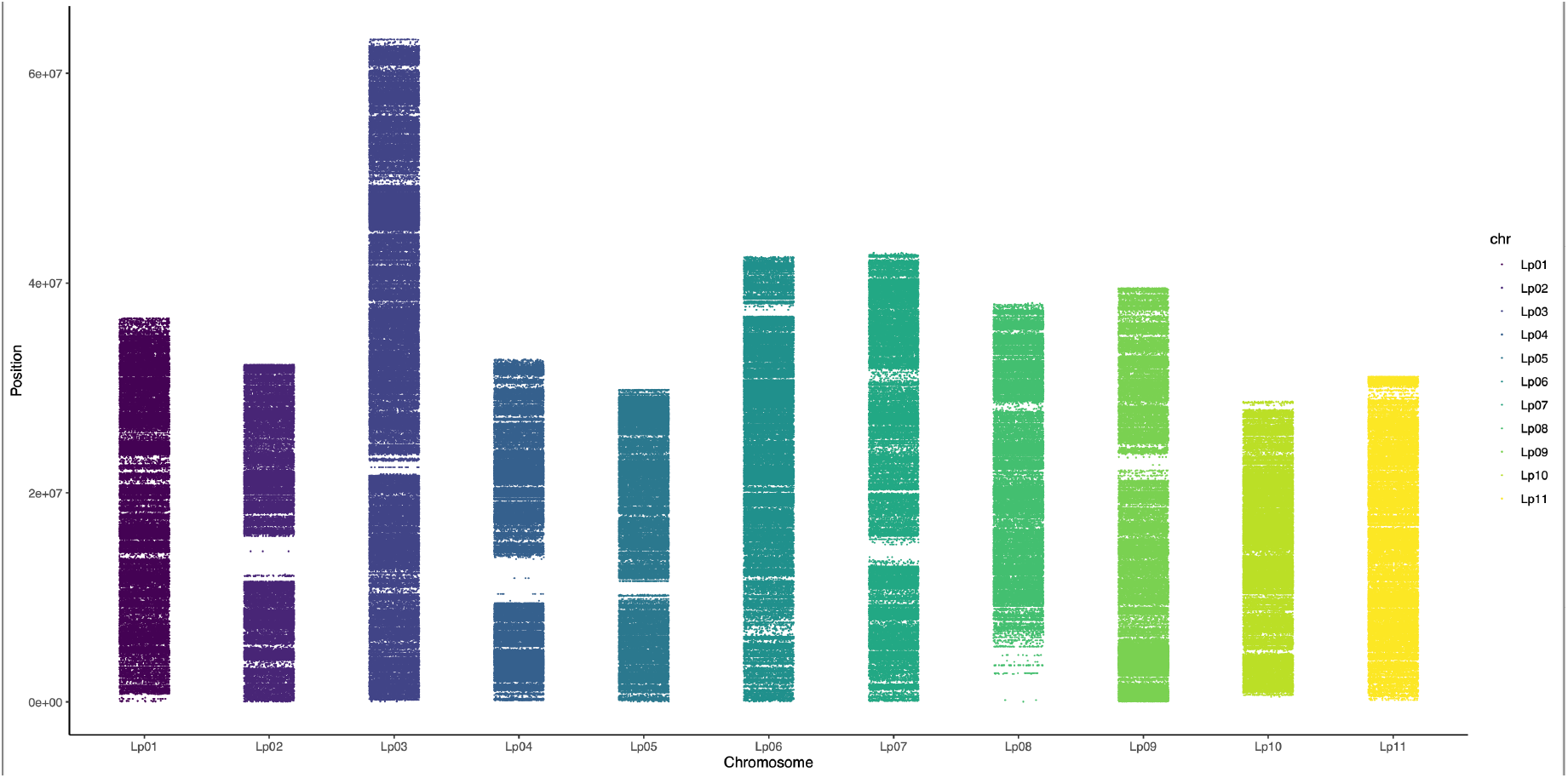
Distribution of highly filtered SNPs (ca. 500k) distributed along the 11 lablab chromosomes

**Supp. Fig 4.**
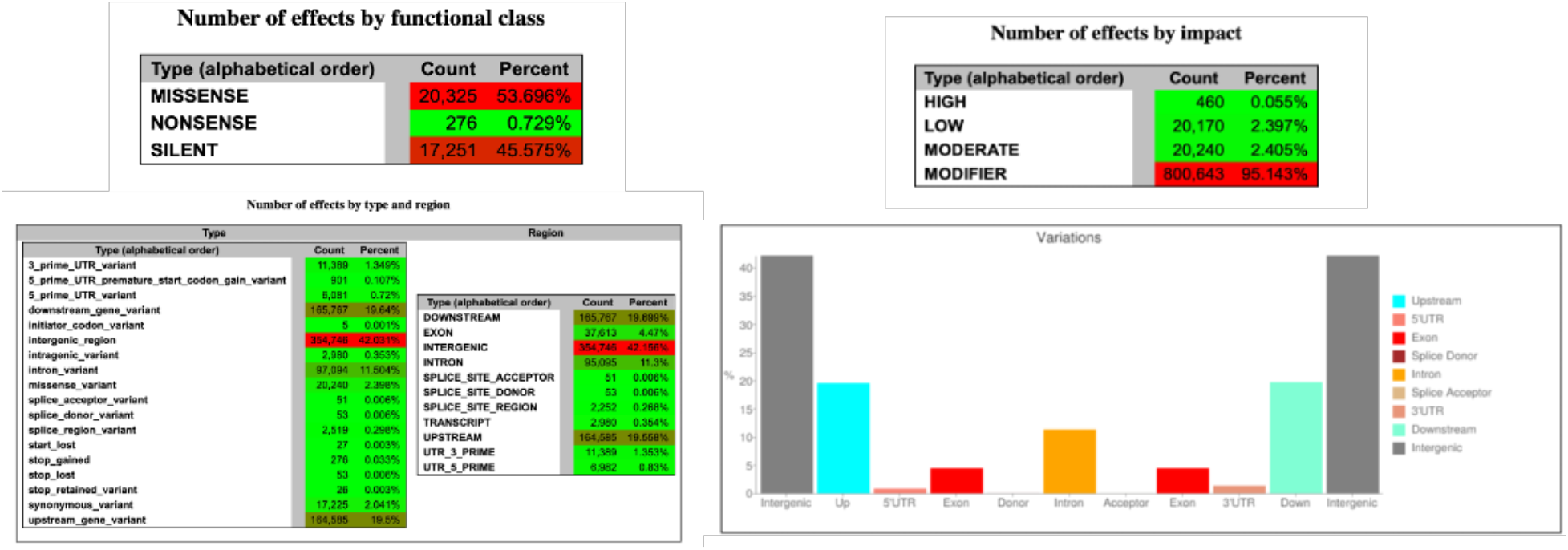
SnpEff analysis of filtered 500k SNPs

**Supp. Fig 5.**
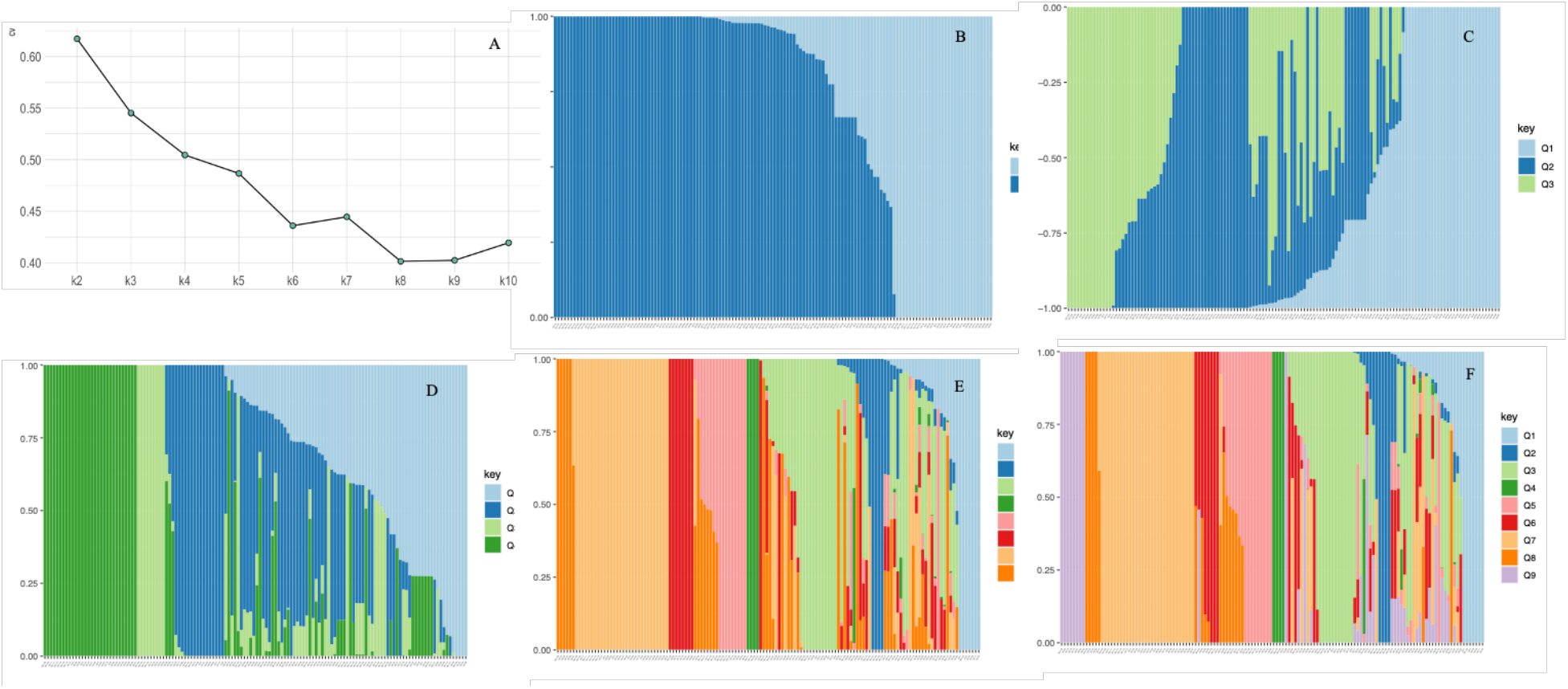
Plot of ADMIXTURE cross validation error from K=2 through 9 (A). Bar plots based on ADMIXTURE analysis, for k=2,3,4,8,9 (B-F).

**Supp. Fig 6.**
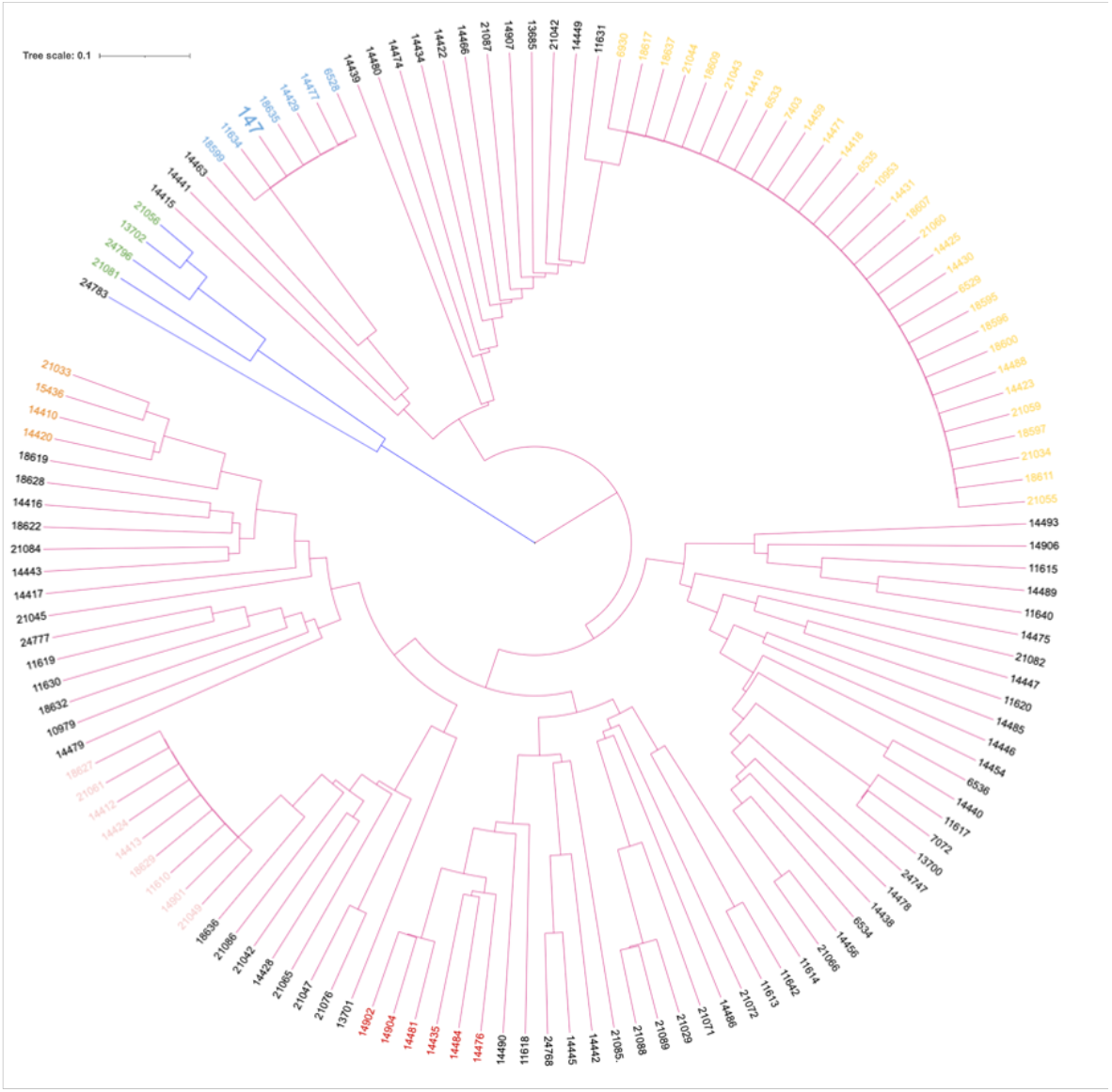
Hierarchical cluster analysis of 136 lablab accessions with ca. half a million genome-wide SNPs. The colored accessions showed single cluster membership in ADMIXTURE analysis. The reference genome, accession 147, is among blue highlighted accessions

**Supp. Fig 7.**
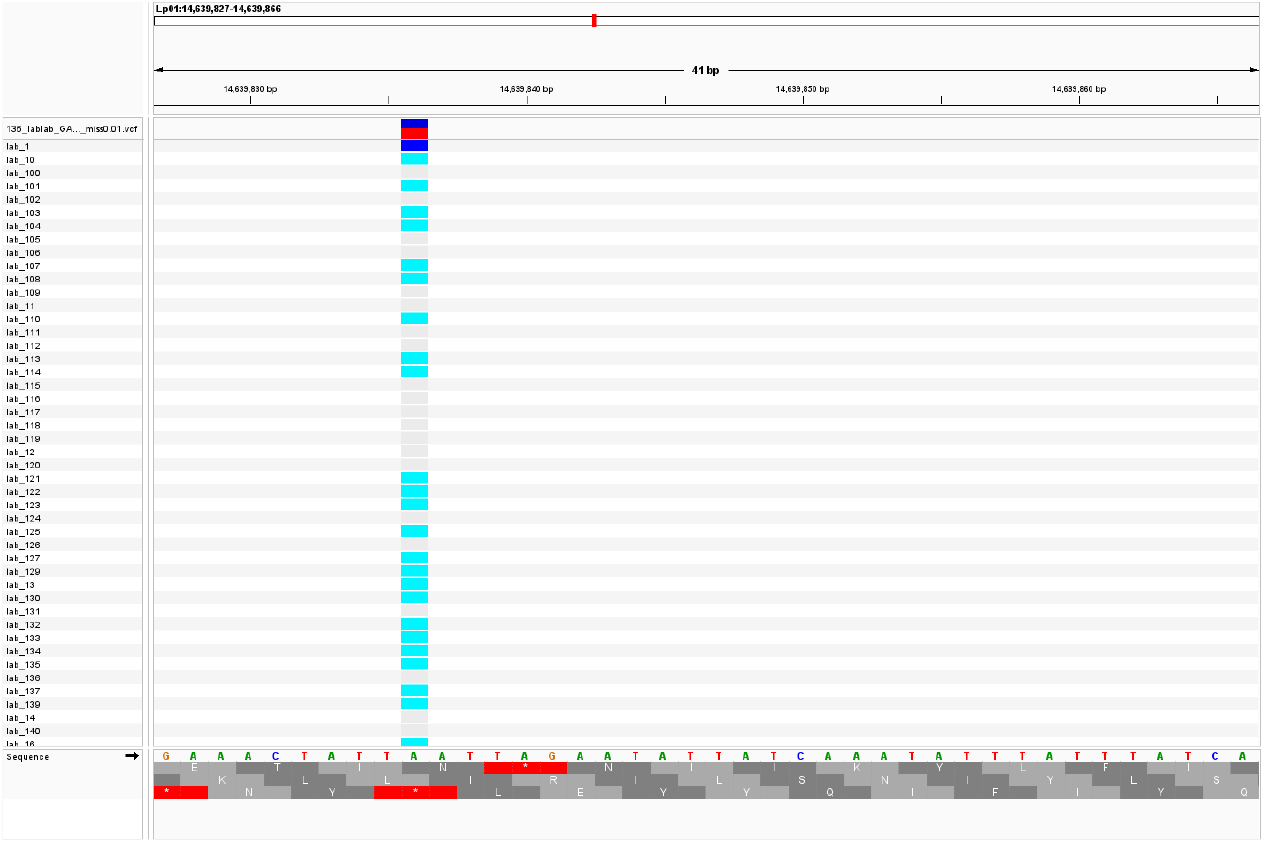
Heatmap of unique/Private SNPs for 127 lablab accessions. Nine accessions from a panel of 136 failed to have private SNPs.

